# Modeling adaptive therapy in non-muscle invasive bladder cancer

**DOI:** 10.1101/826438

**Authors:** Meghan C. Ferrall-Fairbanks, Gregory J. Kimmel, Mik Black, Rafael Bravo, Oana Deac, Pierre Martinez, Maggie Myers, Fereshteh Nazari, Ana Osojnik, Hemachander Subramanian, Yannick Viossat, Freddie Whiting, Roger Li, Karen M. Mann, Philipp M. Altrock

## Abstract

Bladder cancer is the 9^th^ most commonly diagnosed cancer. Nearly half of patients with early stage bladder cancer treated with the immune-stimulating agent BCG have disease recurrence, while 13% progress to invasive bladder cancer. Here we explored the potential of tumor mutational heterogeneity and the role of pro- and anti-inflammatory cytokines to identify different subtypes of bladder cancer that may predict therapeutic response to BCG. Further, we used mathematical modeling of dosing strategies to infer tumor response to varying doses and time schedules f BCG administration. As a proof-of-concept, present adaptive therapy scheduling of BCG as a viable strategy to control tumor size and minimize recurrence.

## I. Introduction

Bladder cancer is the 9th most commonly diagnosed malignancy worldwide, with 81,190 new cases and 17,240 deaths last year in the US [1]. Out of all bladder cancers diagnosed, 75-80% present as non-muscle invasive bladder cancer (NMIBC), a non-aggressive form of the disease which can be managed with therapeutic treatment. NMIBC includes Stage 0 and 1 tumors, while Stages 2-4 are considered muscle invasive (MIBC) invading neighboring muscle and eventually spreading to lymph nodes and beyond [2]. The current standard of care for patients with NMIBC at high risk for progression to MIBC is treatment with intravesical Bacillus Calmette-Guerin (BCG), a vaccine typically given to prevent tuberculosis that has been shown to induce an immune response in bladder cancer patients. BCG is administered in the induction phase weekly for 6 weeks followed by weekly instillations for 3 weeks at specified visits based on cystoscopy/urine cytology results, and once stable, reevaluated every 6 months in the follow-up phase [3]. However, BCG is often not curative; balder cancer recurs in up to 42% of patients treated with BCG and of these patients, 13% will progress to MIBC [3]. The standard treatment for BCG Refractory Disease is surgical removal of the bladder, pelvic lymph node dissection, and urinary diversion.

Currently, there are no biomarkers to predict which patients will recur or advance on BCG. Tumor heterogeneity poses a significant challenge to defining biomarkers[4]. High activation of PI3K signaling in highly proliferative tumors results in poor patient survival in NMIBC and this profile is associated with low immune-response [7, 8]. Furthermore, in MIBC, immune response is associated with better outcomes in patients. Here, we set-out to investigate the relationships between immune response, cellular proliferation, PI3K signaling and recurrence incidence. Further, we used cytokine profiling to model immune phenotypes and patient response to BCG. Finally, we propose several mathematical models as predictive strategies to maximize patient response to BCG while reducing recurrence.

## II. Predicting tumor subclasses using genomic analysis

To establish whether there are molecular markers present in tumors genomes that can be used for subclassfication, we utilized publicly available whole genome expression profiles from 3 datasets: 48 primary pT1 bladder cancers published by Kim et al. [10]; 460 non-muscle-invasive bladder cancers published by Hedegaard et al [7]; and 412 muscle-invasive bladder cancers published by The Cancer Genome Atlas [11]. These data sets were used to investigate the interplay between immune response, cellular proliferation, and PI3K signaling, and their association with altered likelihood of tumor recurrence in bladder cancer. For each of these biological mechanisms (immune response, proliferation, PI3K signaling) singular Value Decomposition was used to generate metagenes from the combined expression profiles of genes known to be involved in these processes (e.g., as previously performed by Nagalla et al. [12]). These expression-based metagenes were then used to identify specific associations with patient survival.

Individually, each of these biological processes were found to be significantly associated with patient prognosis. However, when considered in combination, very specific interactions could be identified. In particular, activation of PI3K signaling was strongly associated with decreased patient survival in highly proliferative tumors, but not in tumors exhibiting lower levels of proliferation. This particular combination (high proliferation and activation of PI3K signaling) was also strongly associated with poor patient outcome, even in the presence of a strong host immune response. In the TCGA data set, higher mutational burden was significantly associated with increased host immune response, however the association between mutational burden and patient prognosis was only present for high proliferation tumors (Fig. 1). A similar relationship has been previously noted in breast cancer [13].

**Figure 1.**
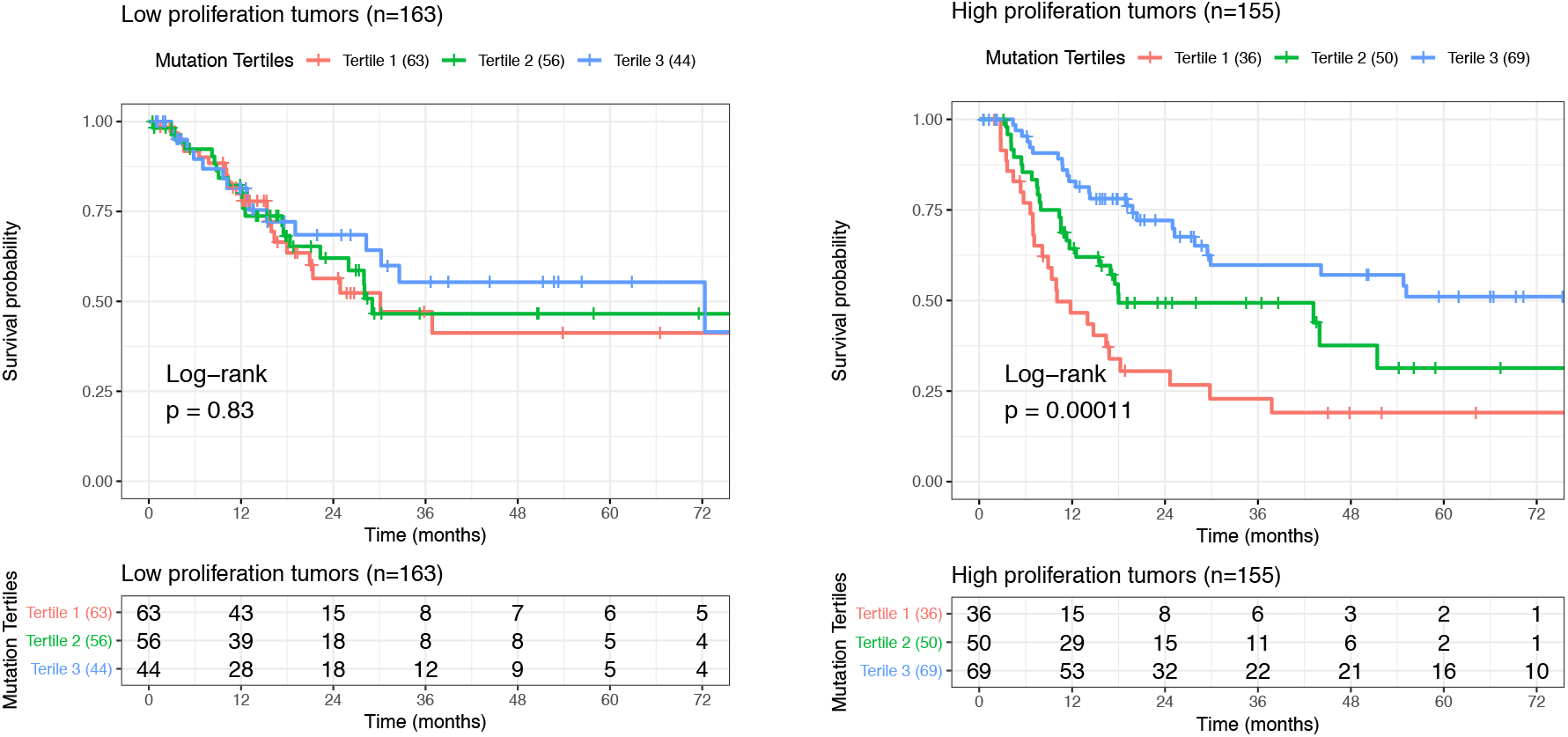
Association between low (tertile 1: 1-112), medium (tertile 2: 113-230) and high (tertile 3: 231-3545) mutation burden, and patient survival. Left: low (below median) proliferation bladder cancers. Right: high (above median) proliferation bladder cancers. Number of patients at risk at each time points are given in the table below each graph.

## III. Modelling approach

### A. Predictive subclasses of bladder cancer

From these genomic analyses, we hypothesized that there are different subclasses of bladder cancer based upon the immunogenicity of the tumor and the status of PI3K activation that will affect patient response to BCG. These two parameters can define four subclasses, illustrated in Figure 2: (1) high-proliferative and non-immunogenic, (2) low-proliferative and immunogenic, (3) high-proliferative and immunogenic, and (4) low-proliferative and non-immunogenic. We predicted that by increasing the immunogenicity of tumors through increasing mutational burden, potentially achieved through radiotherapy, that BCG-refractory tumors will become BCG responsive. Highly proliferative tumors may be targeted by mTOR inhibition, thereby shifting tumors towards a positive BCG-response. To model and quantify the dynamics of these four hypothesized phenotypes, we first explored molecular markers of these phenotypes, then modeled the phenotypic interaction using (1) a deterministic, ordinary differential equations-based approach of the different cancer cell and immune cell populations[9], (2) incorporated contribution of spatial variables in a partial differential equation systems, and (3) explore stochasticity of a bladder tumor microenvironment using an agent-based modeling (ABM) approach. The implementation of these approaches is described in the following subsections.

**Figure 2.**
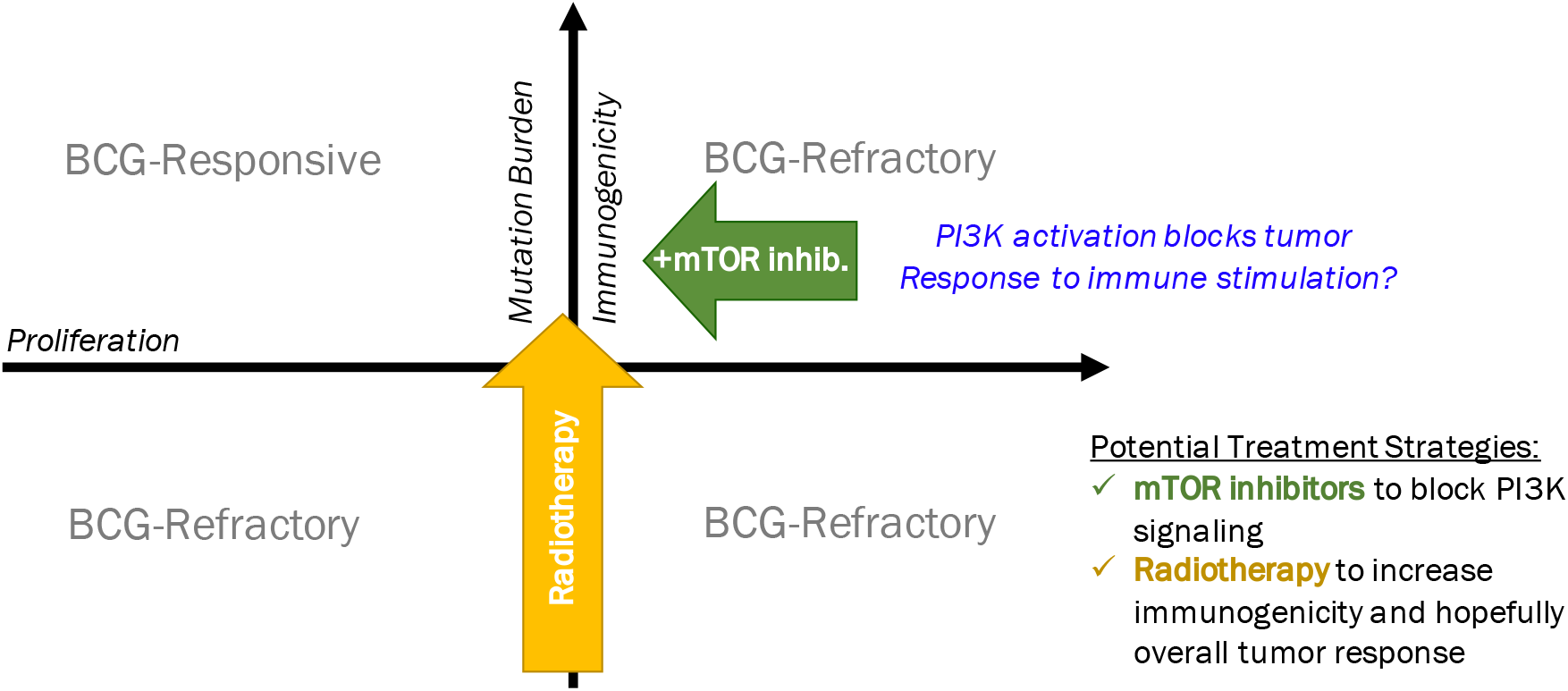
Non-muscle invasive bladder cancer cell behavior can be broadly divided into a proliferative axis an immunogenic axis. Bladder cancer that is immunogenic and low-proliferative responses to BCG-treatment, but as proliferation increases and/or tumor burden decreases, the response to BCG is lost.

### B. Modeling four tumor cell phenotypes of immune invasion response and adaptive therapy scheduling

Here, we employ dynamical systems modeling of phenotypic properties of bladder cancer cell and immune cell response to BCG treatment. A schematic of the bladder cancer immune-phenotypes used to model the tumor-immune cell interactions is depicted in the top panel of Fig 3.

**Figure 3.**
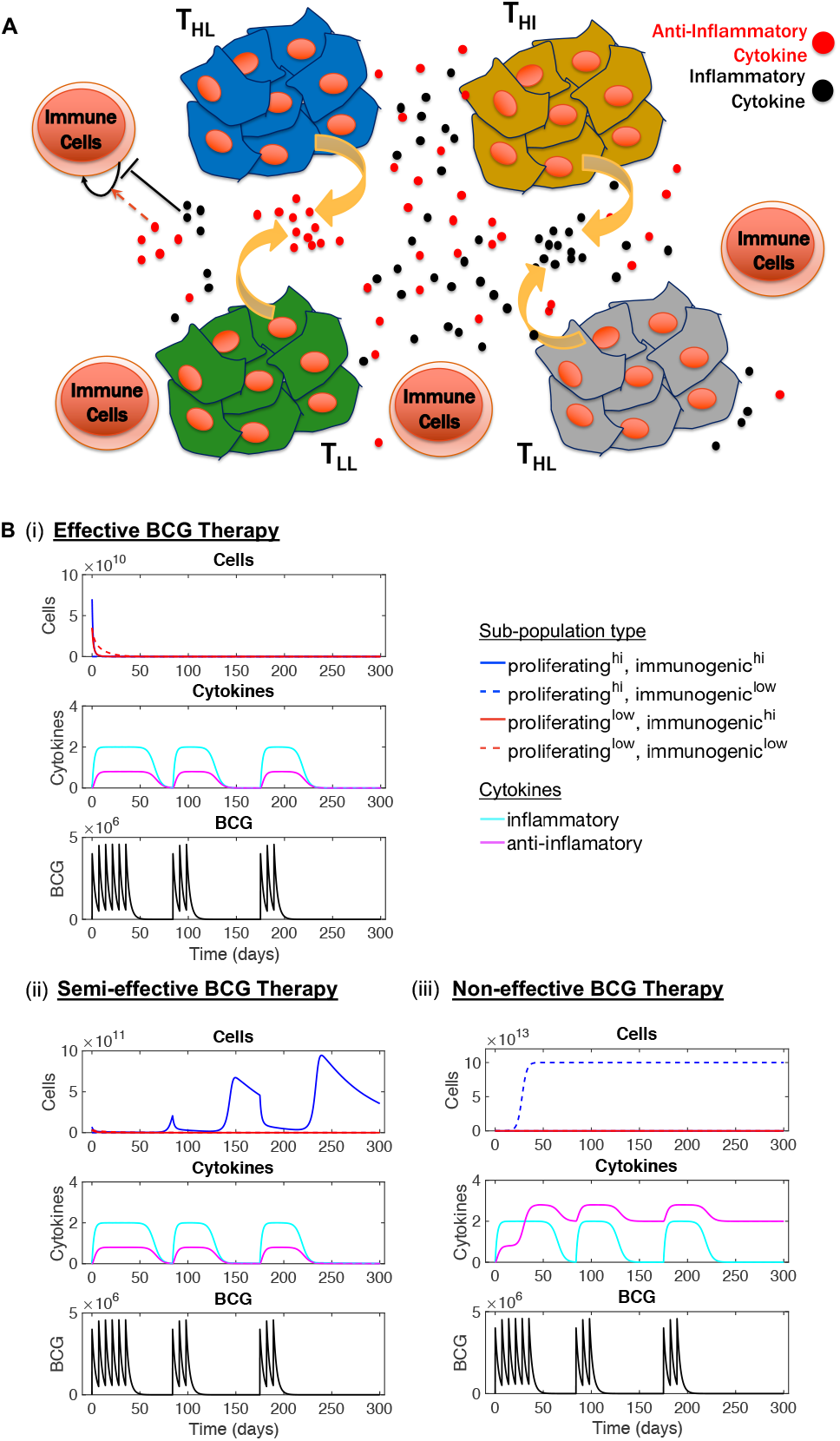
Different immunophenotypes of bladder cancer alter effectiveness of conventional BCG therapeutics. **A**: Schematic of hypothesized bladder cancer cell types and their interactions with secreted cytokines in the tumor microenvironment. **B**: Different tumor compositions and cytokine profiles resulted in different effectiveness of conventional scheduling of BCG administration.

Overall tumor burden (T_B_) is quantified by the sum of the following four different tumor-immuno-phenotypes: high-proliferating, high-immunogenic (T_HI_) bladder cancer cells;high-proliferating, low-immunogenic (T_HL_) bladder cancer cells; low-proliferating, high-immunogenic (T_LI_) bladder cancer cells; and low-proliferating, low-immunogenic (T_LL_) bladder cancer cells. The population dynamics of each of these phenotypes is dependent on the concentration of each of the other phenotypes and the number of inflammatory cytokines (C_I_) and anti-inflammatory cytokines (C_A_), which is regulated by the BCG treatment. The following set of ordinary differential equations describes the population dynamics of the bladder cancer cells driven by density-dependent proliferation and death:

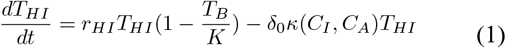

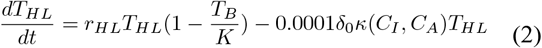

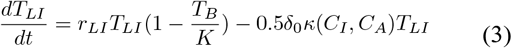

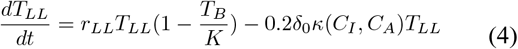

where *r_i_* describes the growth rates of each of the subtypes, *K* is the carrying capacity of the system, and *τ_0_* is the death rate due to the interplay of inflammatory and anti-inflammatory cytokines.

The concentrations of inflammatory and anti-inflammatory cytokines are dependent on BCG treatment. The cytokine concentration is described by the following dynamical system:

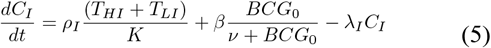

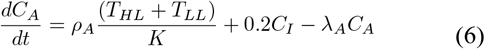

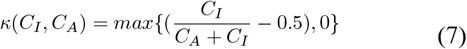

where *ρ_i_* describes the rate of cytokine production, *BCG_0_* is the initial dose of BCG, *β* describes the maximal impact of BCG on inflammatory cytokine production, *ν* describes the effective K_M_ Michaelis-Menten constant of BCG, and *λi* describes the rate of cytokine decay. Modeling conventional BCG treatment outcomes, BCG will lead to tumor killing, if (1) there is increased immune predation, (2) there are more inflammatory cytokines than anti-inflammatory cytokines, and (3) if there are initially no high-proliferating, low-immunogenic bladder cancer cells.

If there are no high-proliferating, low-immunogenic bladder cancer cells and there is a low basal immune predation (*τ_0_*), then the tumor is temporarily kept under control by the traditional BCG treatment, but eventual relapse will occur.

If there are initially high-proliferating, low-immunogenic bladder cancer cells in the initial tumor composition or an immunosuppressive environment with more antiinflammatory cytokines than inflammatory cytokines, there will not be a response to BCG treatment.

However, by using alternative BCG scheduling, specifically shown by reducing time between therapies, we simulated a decreased overall bladder cancer tumor burden, suggesting that adaptive therapy may be more effective than the convention BCG scheduling for at risk individuals who are ultimately unresponsive to BCG therapies (Fig. 4).

**Figure 4.**
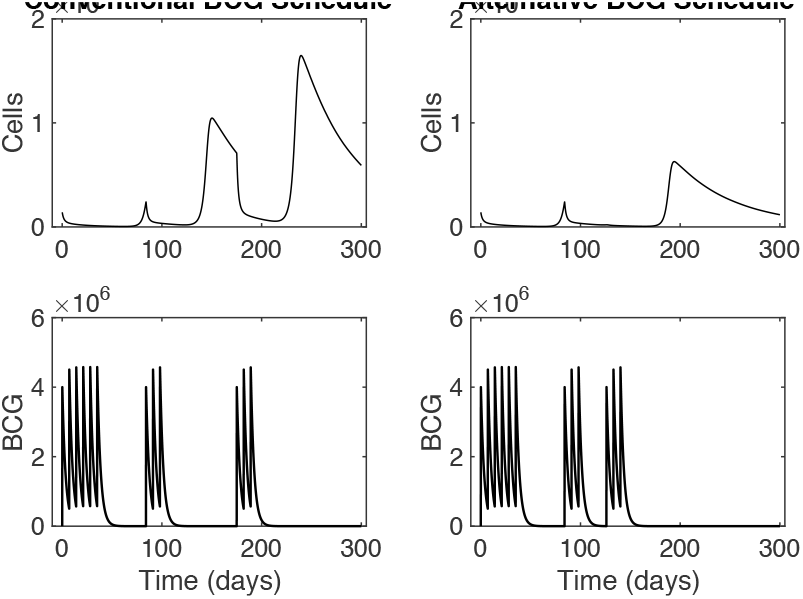
*In silico* perturbations show that intervening early with additional doses of BCG in an adaptive schedule reduced overall tumor budern. This reduction suggests that adaptive BCG scheduling based on tumor composition may improve prognosis of NMIBC patients.

### C. Evolutionary Game Theory of innate immune signaling and adaptive immune cell predation

To model freqeuncy and density dependent dyanmics of interacting cell populations (and drug concentrations), Evolutionary Game Theory provides a promising set of modeing tools[9], highlighted by recent successful modeling approaches[14, 15].

We wanted to explore if surgical researction before BCG administration could be therapeutically beneficial to patients. We hypothesize that on top of eliciting a high, anti-BCG immune response, BCG may infect some tumor cells, potentially leading to exposition of tumor antigens to the immune system (either because infected tumor cells directly expose antigens, or simply because some tumor cells are killed by BCG and tumor debris are exposed to the immune system). To test this hypothesis, we developed a game-theoretic model with three actors: tumor cells, T-cells, and interleukins (IL) (shown in Fig 5). Tumor cells have a natural growth-rate and a death-rate due to the action of T-cells. T-cells production is stimulated by interleukins (more precisely, it is proportional to the product of the number of T-cells and the number of Interleukins). Otherwise the number of T-cells decays exponentially due to a natural death rate. Interleukins production is stimulated when T-cells and tumor cells meet (so is proportional to the product of the number of T-cells and the number of tumor cells), or the concentration of IL decays exponentially. This game-theoretic, differential equations model is thus defined as

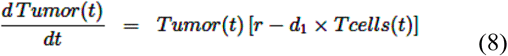

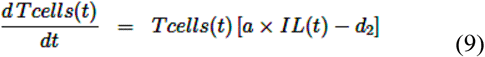

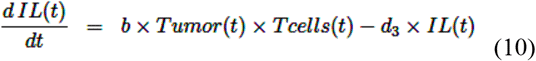

This model could be further augmented by adding saturation of the tumor’s impact on T-cells: for very large tumors, the death rate of tumor cells induced by the presence of T cells becomes less than proportional to the number of T cells. This would allow the tumor to escape the immune system, if it grew very large.

**Figure 5.**
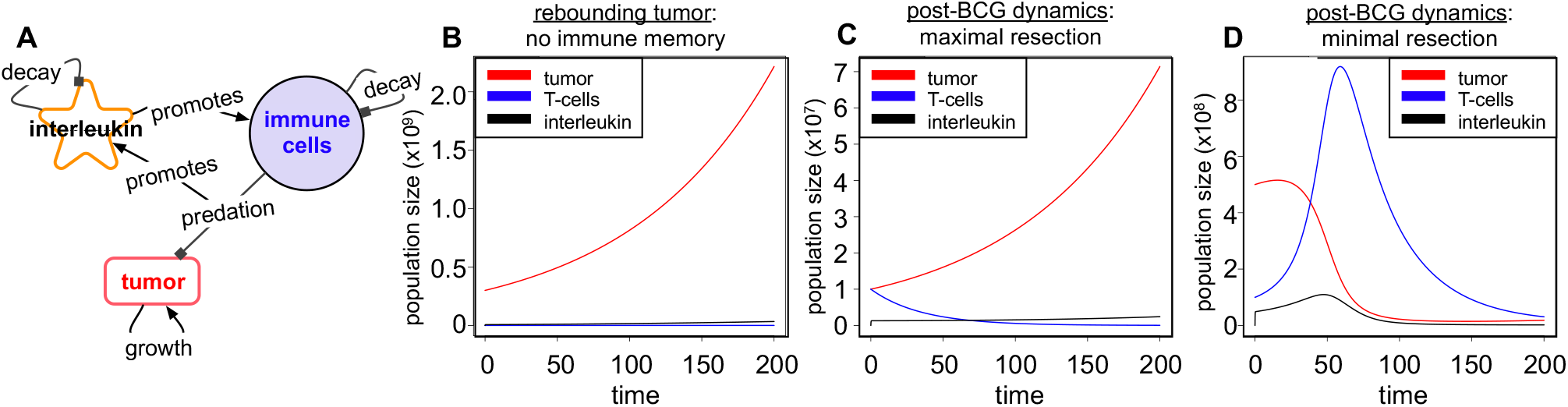
Game-theoretic model exploring full or partial surgical resection of non-muscle invasive bladder cancer prior to current BCG therapeutic dose schdeuling. **A:** Model schematic of a general game-theoretic approach that includes tumor, T cells, and immune-stimulants (interleukins). **B:** Dynamics following maximally possible surgical resection. **C:** Dynamics following resection and immune memory. **D:** Resection and BCG.

We used this modeling approach to address the following question: If the tumor burden is higher that a particular threshold, it may indeed activate an increased immune response, but what impact does such dynamic feedback have in the long run? Could it be of benefit to not aim for complete tumor eradication simply due to this feedback? The dynamics of eqs. (8)–(10) show that the long-term behavior may indeed be improved by leaving some tumor after initial treatment/surgery, see Fig. 5.

### D. Mutation load (immunogenicity) vs. proliferation rate increases

For many phenotypes, the difference in trait and genotype space (Fig. 1) may be minimal. In this case we can move from a coupled model of ODEs towards a continuous description, where the phenotypes vary in a continuous fashion along two major dimensions—mutation burden and proliferation rate. Change in this two-dimensional space over time is then a result of tumor evolution and of response to (BCG) treatment. We here define the cell concentration *c*, a normalized mutational load *m* ∈ [0,1], and a basal proliferation rate *r*. The phenotype-specific birth rate can then be defined by equation (14):

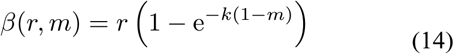

Where we have introduced the parameter *k*, a measure of the influence mutational burden has on growth rates. Similarly, we introduce the basal death rate *ρ*, the phenotype-specific death rate can be defined by equation (15):

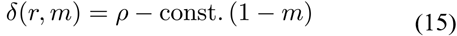

The constant controls the sensitivity of mutational load on the death rate, where we assume high mutational load is the most sensitive and low mutational load is the least sensitive.

Assuming the number of phenotypes is large we can move from discrete compartments to a continuous range in trait space. This notion defines a partial differential equation of how the change in cell concentration changes over time in trait space (r, m) and is given by:

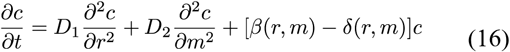

where D_1_ is the rate of phenotype variation in the growth rate, D^2^ is the mutational rate, and 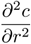 denotes the partial derivative with respect to baseline proliferation rate.

We observed shifts in high concentration of cells in (r, m) space that depend on whether the treatment is on or not (Fig. 6); as an effect of mutation burden-associated immunogenicity, treatment penalizes highly proliferating, highly mutagenic cells the most. This is consistent with the migration of phenotypes around trait space in this model. In the absence of treatment, we saw shifts toward a majority of high proliferating cells. When treatment was turned on, we saw an initial decrease in total cancer cell number, as the cell population moved towards a more moderate proliferation rate distribution, as faster proliferating cells die during treatment.

**Figure 6.**
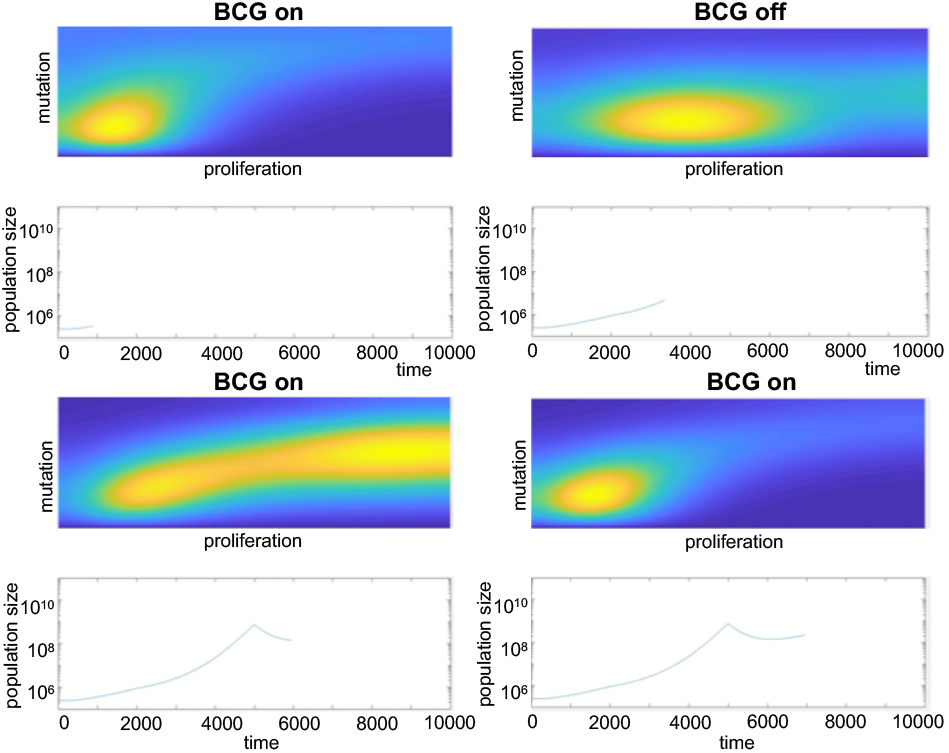
Movement in phenotype-space based on whether treatment is on or off. With BCG therapy on, the cancer cell population favors low proliferative and mutational burden; however, while it is turned off, the highly proliferative subtype begins to thrive. We observe initial tumor control while the cancer phenotype adjusts to its changing environment, but eventually it continues to grow. This suggests modulating the dose and therapy time to keep the “sensitive” cells competing with the “resistent” cells.

**Figure 7.**
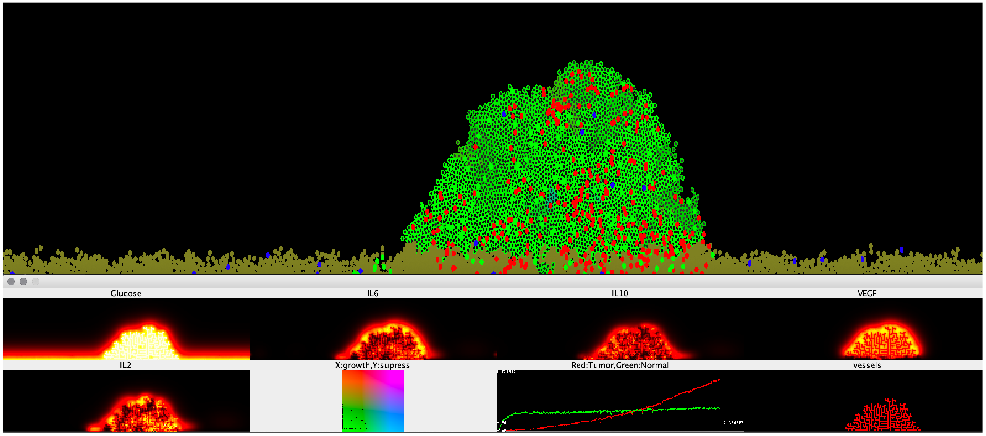
Bladder epithelium as a hybrid ABM model visualization. Top: Off-Lattice ABM, brown cells are normal epithelium, green cells with black centers are tumor cells (variations in cell color correspond to phenotypes), red cells are effector T Cells, blue cells are naïve T Cells, and green cells with no centers are anergic T Cells. Bottom: PDE concentrations, a map of phenotypes, populations over time, and vasculature are shown. Downward spikes in the population plot show where BCG therapy was applied.

### E. Hybrid Agent Based Model of Bladder Epithelium

To explore the effects of local interactions on overall outcome, and to combine several modeling assumptions stated above, we built a hybrid ABM of the homeostatic bladder epithelium with blood vessels and immune cell surveillance. This served as an alternative substrate to model tumor evolution and the effects of treatment. Cells are modeled off-lattice and grow to a height limited by glucose availability (normally around 6 cell diameters). Tumor cells are able to circumvent this due to release of VEGF, which stimulates vessel growth. Vessels are modeled with an on-lattice ABM. In high VEGF tumors, these rules will lead to polyp formation, in low VEGF tumors, cells will instead tend to compete more with the epithelium and grow in a sheet.

The tumor cells start with a phenotype that is identical to the normal tissue, except that they can produce VEGF and mutate. Mutation can occur along two axes: a growth axis, which increases proliferation rate, and an immunosuppression axis, which makes T cell anergy more likely, and a proliferation axis, which increases proliferation rate. The cost of either type of mutation is that it increases the cell’s antigenicity, which increases the chance of immune detection. If immune predation is low, cells will specialize in rapid proliferation to outcompete other clones. If immune predation is high, cells will specialize in immunosuppression first to prevent mutations from leading to cell killing, then become rapidly proliferating once they are sufficiently immunosuppressive.

The model contains IL2, IL6, and IL10 cytokine PDEs, which mediate the immune response. IL2 and IL6 are pro-inflammatory. IL6 is released upon tumor cell death to represent tumor antigen release. IL-6 and IL-2 are both released by effector T cells, while IL-10 is released by anergic T cells.

The immune cell presence in the model is currently limited to just CD-8 T cells. These cells enter the domain through the vasculature proportional to the local IL6 and IL2 concentration, with a penalty for the local IL10 concentration. T cells in the model can exist in 3 possible states: naïve, effector, and anergic. When cells first enter through the vasculature, they are in the naïve state. upon identifying an antigenic tumor cell, these cells can switch to either the effector or anergic state depending on the local concentration of cytokines and the immuno-suppressiveness of the tumor cells. Aging effector T-cells will automatically become anergic, dampening the immune response over time.

This spatially explicit model of bladder cancer ecology and evolution can also incorporate BCG therapy. BCG causes all cells in the model to become “infected” making them highly immunogenic. This change in immunogenicity causes a transient strong cytotoxic response which is eventually dampened as the T-Cells become exhausted. Multiple spaced pulses of BCG can constrain tumor growth and even cure in more benign cases. Future efforts should involve further calibration, testing other dosing schedules, and incorporating other immunotherapies such as anti-PD/PDL-1 to prevent immunosuppression.

## IV. Discussion

Of patients diagnosed with non-muscle invasive bladder cancer, 30-40% are unresponsive to BCG, necessitating life-altering surgical intervention. We have shown a relationship between mutation burden and patient survival, specifically in tumors that exhibit high expression of genes driving cellular proliferation. PI3K activation identified a relationship between PI3K activation, poor immune response to BCG, and a concomitant decreased patient survival.

Here we have presented multiple different modeling approaches that seek to build upon these observations and, using complementary subsets of biological assumptions, predict new adaptive approaches to treating bladder cancer patients. We hypothesized that there are multiple ways in which currently-available therapeutic options can be used in *combination* with BCG, or in *alternative scheduling,* to improve response to treatment. First, we modeled how PI3K activation abrogates a patient’s response to BCG and is found in high-proliferative and high-immunogenic (tumor burden) tumors. We postulate the use of current mTOR inhibitors to shift the tumor along the proliferative axis (to the left in Fig. 2) and make the tumor more susceptible to immune-therapeutics, such as BCG. Next, we modeled how tumor resection prior to therapy could be improved toward stronger immune-stimulating signaling and stimulation by additional therapy in order to cause long-lasting treatment benefits. Finally, we modeled the effects of boosting immunogenicity of BCG-refractory bladder cancers on improved patient responses to treatment. We postulate that by utilizing radiotherapy on bladder cancer, clinicians will shift the tumor along the immunogeneic/mutational burden axis (shifting up in Fig. 1) and increase the immune response to BCG by increasing the mutation burden and increasing the cell killing and release of cytokines and antigens prior to BCG treatment. Taken together, our models suggest that adaptive BCG therapy based on patient-specific tumor composition would enable clinicians to exert improved control of tumor burden and improve patient outcomes.

## V. Data Availability

Code associated with the ODE, EGT, and PDE models is available at: https://github.com/mcfefa/2018-IMO-Wksp-TeamRed

## About the Authors

M.C.F., G.J.K., R.B., and P.M.A. are with the Department of Integrated Mathematical Oncology, H. Lee Moffitt Cancer Center and Research Institute, Tampa, Florida, USA.

M.B. is with the Department of Biochemistry, University of Otago, Dunedin, New Zealand.

O. D. is with St. James’s Hospital Dublin, County Dublin, Ireland.

P. M. is with the Université Claude Bernard Lyon 1, Centre Léon Bérard, Cancer Research Center of Lyon, Lyon, France

M.M. is with St. Jude Children’s Research Hospital, Memphis, Tennessee, USA.

F.N. is with the University of Chicago, Chicago, Illinois, USA.

A.O. is with the Mathematical Institute, University of Oxford, Oxford, United Kingdom.

H.S. is with Department of Cancer Physiology, Moffitt Cancer Center and Research Institute, Tampa, Florida, USA.

Y.V. is with the Department of Mathematics, Université Paris-Dauphine, Paris, France.

F.W. is with Department of Oncology and Cancer Biology, Barts Cancer Institute, London, United Kingdom. (CHECK)

R.L. is with the Department of Genitourinary Oncology, H. Lee Moffitt Cancer Center and Research Institute, Tampa, Florida, USA.

K.M.M. is with Departments of Molecular Oncology, Gastrointestinal Oncology, and Malignant Hematology, H. Lee Moffitt Cancer Center and Research Institute, Tampa, Florida, USA and Department of Oncological Sciences, College of Medicine, University of South Florida, Tampa, Florida, USA.

## Acknowledgment

We would like to thank the Moffitt Cancer Center Department of Integrated Mathematical Oncology (IMO) Chair, Dr. Alexander Anderson, for organizing the 8^th^ Annual Moffitt IMO Workshop: Evolutionary Therapy, where this project was conceived. We are also extremely grateful for Moffitt Cancer Center and the Moffitt PSOC for support, through NCI grant U54CA193489.

